# Exploring taxonomic diversity in the genus *Suberites* Nardo, 1833: Insights from microbial, phylogenetic, and morphological analyses

**DOI:** 10.64898/2026.11.12.765623

**Authors:** Verónica Llorente-Vaquero, Carlota Gracia-Sancha, Francisco Baldó, Marta Turón, Marta Miñarro, Andrea Corral-Lou, Robert Cassidy, Javier Cristobo, Sergi Taboada, Ana Riesgo

## Abstract

Marine sponges (phylum Porifera) are a highly diverse and polymorphic group of marine invertebrates whose taxonomic delimitation is often challenging due to high intraspecific variability and the scarcity of diagnostic morphological characters. Their close association with microbial symbionts also makes them an important source of secondary metabolites, some of which may contribute to sponge colouration. Here, we use morphological characters, molecular phylogenetic markers, and 16S metabarcoding of the symbiont community to understand speciation in a common Mediterranean sponge lineage. We collected 43 specimens of the genus *Suberites* from the Mediterranean Sea and the North and South Atlantic Oceans, between 55– 150 m depths, and sequenced two barcoding genes, cytochrome c oxidase I and 28S rDNA, as well as studied their spicular complements. We identified five closely-related species: *S. domuncula*, *S. ficus/pagurorum*, *S. dandelenae*, *S. ambulodomus*, and a new species of *Suberites* from the Balearic Islands, *Suberites* sp. nov. The latter represents a potential new pseudo-cryptic species closely related to *S. domuncula*, because of their strong morphological and molecular affinities. The potential new Mediterranean species exhibits six colour morphotypes (yellow, gray, orange, brown, red, and blue), and the microbiome analyses of the symbiotic communities revealed significant differences in bacterial community composition among colour morphotypes, specially of bacteria that produce different pigments which potentially give the sponge is colouration. These results provide valuable insights into species boundaries and evolutionary relationships within the genus *Suberites* and support a role for the microbiome in sponge colouration.

## INTRODUCTION

Sponges (phylum Porifera) are one of the oldest extant groups of metazoans (Rossi et al., 2026), which positions them as a highly successful lineage, largely due to their ability to adapt to their environment, allowing them to colonize almost all ecosystems in both marine and freshwater habitats (van Soest et al., 2021). Traditionally, taxonomic identification within this phylum has relied on morphological characters, including external features such as body form, colour, and texture (Hooper & van Soest 2002), as well as internal characteristics, in which skeletal architecture and spicule morphology, including type, shape, size, and dimensions, constitute key taxonomic traits (Cárdenas et al., 2007). However, morphological plasticity, overlapping diagnostic characters, and the presence of cryptic species can complicate sponge identification and classification (Xavier et al., 2010). This taxonomic ambiguity is particularly acute when evaluating external phenotypic traits such as colour (Pankey et al., 2024). Living colouration is notoriously diverse across demosponges, exhibiting striking polychromatism where single species encompass multiple contrasting colour morphs (Becking et al., 2013; Guiollot et al., 2026). To overcome these limitations, integrative taxonomy has emerged as a powerful approach for accurately delimiting species, combining traditional morphological analyses with molecular data (Swierts et al., 2017; Santodomingo et al., 2024).

The genus *Suberites* (Nardo, 1833) encompasses 88 described species (de Voogd et al., 2026) distributed globally with an apparently cosmopolitan character, occurring frequently in cold and temperate marine waters (van Soest, 2002; van Soest et al., 2008; Turner et al., 2024; Payne et al., 2024). Across its bathymetric gradient, the members of this genus occupy settings ranging from the low intertidal and shallow subtidal kelp forests between 0 – 30 m down to outer continental shelf and bathyal depths exceeding 180 - 242 m and into deep subpolar waters (Austin et al., 2014; Turner et al., 2024; Payne et al., 2024). Ecologically, *Suberites* colonizes both hard rocky substrates and unconsolidated soft sediments; in soft-bottom environments, species exhibit specialized adaptations such as basal anchoring stalks or facultative mutualisms with hermit crabs (e.g., *Pagurus liochele* (Barnard, 1974), *Pagurus bernhardus* (Linnaeus, 1758)) (van Soest, 2002; Payne et al., 2024). Morphologically, the genus is characterized by an ectosomal barrier of smaller tylostyles over a choanosomal reticulation composed of larger megascleres, dominated by tylostyles and occasionally smooth or spined centrotylote microstrongyles (van Soest, 2002). In living specimens, the coloration recorded across *Suberites* species spans vivid orange, blue, reddish-brown, yellow, brownish-yellow, pale cream, and mottled beige marked with dark gray or black patterns (Connes et al., 1974; Jaklin et al., 2007; Becking et al., 2009; Payne et al., 2024). While live coloration can provide useful field cues at regional scales, its diagnostic utility remains fundamentally limited, as the same species encompassed very different colorations patterns (Becking et al., 2009; Austin et al., 2014; Turner et al., 2024). Furthermore, sponge coloration is highly susceptible to external and internal influences, including the qualitative profile and concentration of cellular pigments (such as carotenoids and melanin), the depth and ambient light regime of the habitat, surface fouling and trapped fine sediment particles, interactions with microbial or algal endosymbionts, and transient physiological or reproductive states (López-Legentil et al., 2010; Becking et al., 2013; Maia et al., 2021). However, the scarcity of discrete diagnostic characters alongside pronounced morphological plasticity (spanning colouration, growth forms from encrusting to massive or stalked, as well as wide variation in spicule dimensions, head morphologies, and microsclere abundance) hinders specimen identification in this genus (van Soest, 2002; Austin et al., 2014; Turner et al., 2024; Payne et al., 2024). Such extensive character overlap between distinct lineages generates substantial obstacles for the recognition of stable morphospecies (Becking et al., 2013; Samaai et al., 2017). Consequently, morphologically defined taxa have historically masked complexes of cryptic or pseudocryptic species, which increasingly require multi-locus molecular sequencing (*cox1*, *28S* rDNA) for evolutionary resolution (Solé-Cava & Thorpe, 1986; Samaai et al., 2017).

Sponges have strong symbiotic associations with bacteria and archaea, which supply them with additional metabolic and defensive properties (Taylor et al., 2007; Hentschel et al., 2012). Symbiotic microorganisms can directly shape sponge coloration either by synthesizing pigments *de novo* or by altering host metabolic pathways (Erwin & Thacker, 2007; Hentschel et al., 2012). In well-illuminated shallow environments, phototrophic symbionts such as cyanobacteria often serve as the primary drivers of tissue pigmentation, conferring distinctive hues and photosynthetically derived nutrition to the host (Erwin & Thacker, 2007: Usher 2008). Conversely, in mesophotic and deeper shelf zones where species of *Suberites* frequently occur, the attenuation of incident light drastically limits phototrophic assemblages, leaving the ecological mechanisms and potential microbial contributions to pigmentation far less understood (Idan et al., 2018; Olson et al., 2010). While characteristic coloration in deeper demosponges has commonly been attributed to endogenous carotenoid complexes, whether non-photosynthetic heterotrophic bacteria correlate with or modulate these pigments remains largely untested (Sacristan-Soriano et al., 2019; Moreno-Giménez 2026). A well-known case is *Suberites domuncula* (Olivi, 1792), which displays wide chromatic variation (yellow, brown, white, orange-red, and bluish-red) and whose associated microbiota has long been suspected to play a role in pigmentation (Cariello and Zanetti, 1981; Jaklin et al., 2007; Müller et al., 2012). This polychromatism presents substantial taxonomic challenges, as distinct color morphs may easily be mistaken for separate morphospecies or mask cryptic taxa unless evaluated through an integrative approach combining morphology and multi-locus molecular sequencing (Solé-Cava & Thorpe, 1986; Samaai et al., 2017; Turner et al., 2024;). Consequently, an important knowledge gap remains regarding whether the distinct colour patterns observed in *Suberites* are associated with differences in microbiome composition, host genetic identity, or environmental regimes (Becking et al., 2013; Turner et al., 2024; Morozov et al., 2025).

For this study, *Suberites* specimens displaying diverse chromatic morphotypes (including gray, yellow, orange, red, brown, and blue) were collected from mesophotic depths around the Balearic Islands in the western Mediterranean Sea and were assessed alongside comparative congeneric material from the Adriatic Sea and the east Atlantic. Addressing this system targets three interconnected knowledge gaps: first, species delimitation within *Suberites* remains notoriously challenging due to extensive morphological plasticity and overlapping skeletal traits; second, phylogenetic relationships and true lineage diversity across the family require more comprehensive multigene resolution to untangle polyphyletic groupings and unmask cryptic complexes; and third, whether these prominent chromatic morphotypes correlate with shifts in symbiotic microbiome composition, localized environmental factors, or host evolutionary divergence represents an open question. Consequently, this study combines morphological examination, molecular phylogenetics, and microbiome profiling to evaluate regional diversity in the genus *Suberites* and explore the biological determinants underpinning their pronounced phenotypic variation.

## MATERIALS AND METHODS

### Sample collection and fixation

We collected a total of 43 specimens of the genus *Suberites*, including *Suberites ambulodomus* (n = 3), *Suberites dandelenae* (n = 3), *Suberites domuncula* (n = 2), *Suberites ficus/pagurorum* (n = 10), and *Suberites* sp. nov. (n = 25), belonging to five different areas: Croatia, Ireland, Spain (Balearic Islands and Galicia), and Namibia (Figure 1 and Table 1).

**Figure 1.**
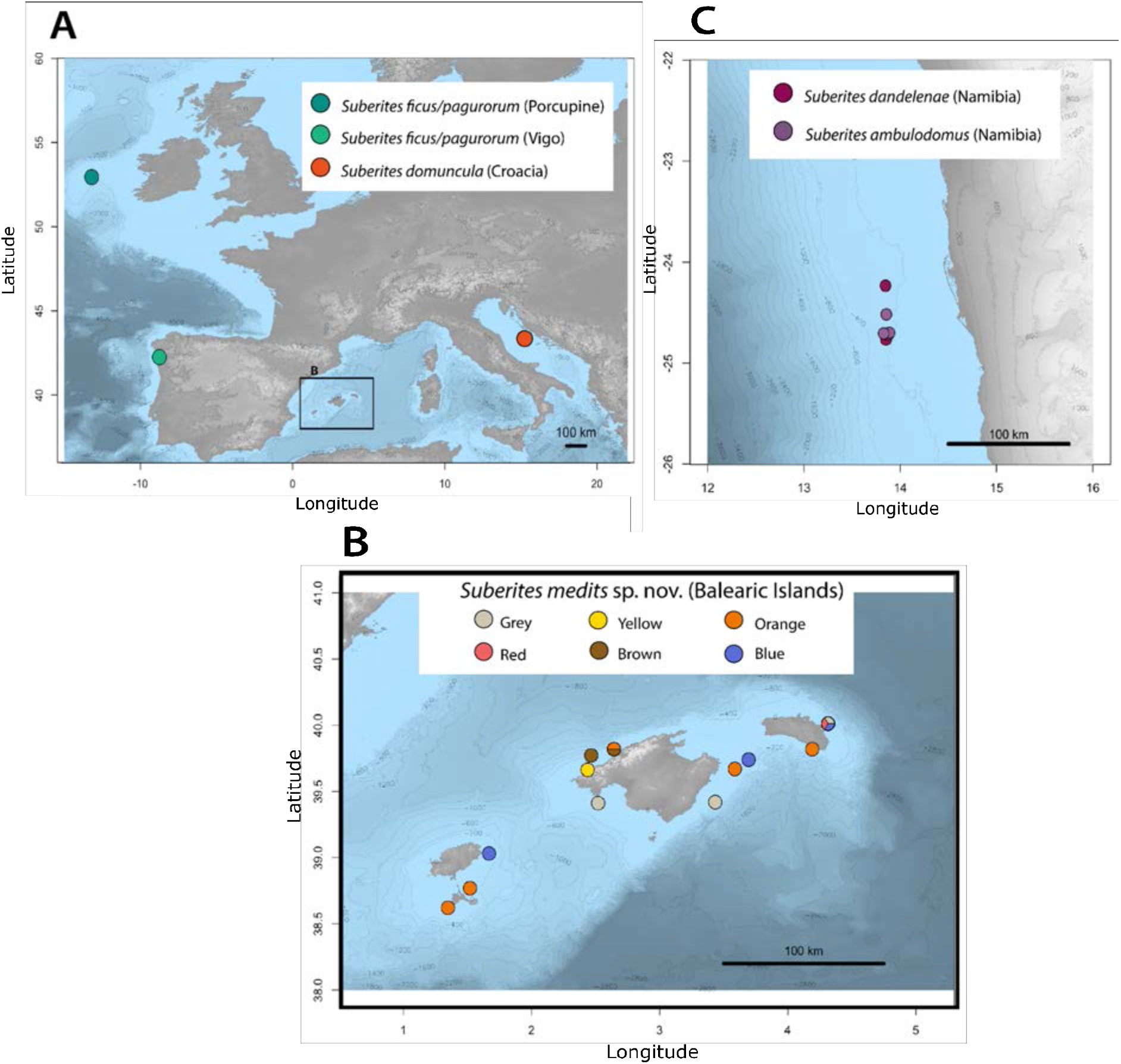
Maps of the sampling areas (exact locations are provided in Table 1). A. Map of Europe showing the sampling sites in the Northern Hemisphere. B. Enlarged map of the European sampling area in the Balearic Islands, showing the sampling sites and the color morphs present at each site. C. Map showing the sampling sites along the coast of Namibia, Africa.

**Table 1.** Data for the samples analyzed in this study. Sample code, identified species, color, sampling location with the corresponding coordinates and depth, and date (unknown for some samples).

| SAMPLE | SPECIES | COLOUR | LOCATION | LATITUDE | LONGITUDE | DEPTH (m) | COLLECTION DATE |
|---|---|---|---|---|---|---|---|
| MED201-2 | *Suberites* sp.nov | Grey | Balearic Islands | 39.41175 | 2.5189 | 69 | 14/6/24 |
| MED217-2 | *Suberites* sp.nov | Grey | Balearic Islands | 39.41935 | 3.43355 | 75 | 18/6/24 |
| MED249-8-3 | *Suberites* sp.nov | Grey | Balearic Islands | 40.0122 | 4.31425 | 78 | 27/6/24 |
| MED249-8-6 | *Suberites* sp.nov | Grey | Balearic Islands | 40.0122 | 4.31425 | 78 | 27/6/24 |
| MED249-8-7 | *Suberites* sp.nov | Grey | Balearic Islands | 40.0122 | 4.31425 | 78 | 27/6/24 |
| MED204-16 | *Suberites* sp.nov | Yellow | Balearic Islands | 39.6636 | 2.438 | 67 | 15/6/24 |
| MED204-21 | *Suberites* sp.nov | Yellow | Balearic Islands | 39.6636 | 2.438 | 67 | 15/6/24 |
| MED224-1-2 | *Suberites* sp.nov | Orange | Balearic Islands | 39.8195 | 4.19165 | 56 | 20/6/24 |
| MED226-5-3 | *Suberites* sp.nov | Orange | Balearic Islands | 39.67035 | 3.58675 | 71 | 21/6/24 |
| MED281-7-2 | *Suberites* sp.nov | Orange | Balearic Islands | 38.6215 | 1.3469 | 58 | 17/8/24 |
| MED282-10 | *Suberites* sp.nov | Orange | Balearic Islands | 38.76945 | 1.51925 | 63 | 18/8/24 |
| MED197-4 | *Suberites* sp.nov | Brown | Balearic Islands | 39.81925 | 2.643 | 69 | 13/6/24 |
| MED197-2 | *Suberites* sp.nov | Brown | Balearic Islands | 39.81925 | 2.643 | 69 | 13/6/24 |
| MED198-1 | *Suberites* sp.nov | Brown | Balearic Islands | 39.77235 | 2.46515 | 146 | 13/6/24 |
| MED249-9 | *Suberites* sp.nov | Red | Balearic Islands | 40.0122 | 4.31425 | 78 | 27/6/24 |
| MED249-9-10 | *Suberites* sp.nov | Red | Balearic Islands | 40.0122 | 4.31425 | 78 | 27/6/24 |
| MED249-9-2 | *Suberites* sp.nov | Red | Balearic Islands | 40.0122 | 4.31425 | 78 | 27/6/24 |
| MED249-9-3 | *Suberites* sp.nov | Red | Balearic Islands | 40.0122 | 4.31425 | 78 | 27/6/24 |
| MED249-9-4 | *Suberites* sp.nov | Red | Balearic Islands | 40.0122 | 4.31425 | 78 | 27/6/24 |
| MED249-9-5 | *Suberites* sp.nov | Red | Balearic Islands | 40.0122 | 4.31425 | 78 | 27/6/24 |
| MED227-5 | *Suberites* sp.nov | Blue | Balearic Islands | 39.73995 | 3.6946 | 81 | 21/6/24 |
| MED227-5-3 | *Suberites* sp.nov | Blue | Balearic Islands | 39.73995 | 3.6946 | 81 | 21/6/24 |
| MED249-11-2 | *Suberites* sp.nov | Blue | Balearic Islands | 40.0122 | 4.31425 | 78 | 27/6/24 |
| MED249-11-3 | *Suberites* sp.nov | Blue | Balearic Islands | 40.0122 | 4.31425 | 78 | 27/6/24 |
| MED292-15 | *Suberites* sp.nov | Blue | Balearic Islands | 39.0315 | 1.66765 | 75 | 1/1/00 |
| NAM56d | *Suberites ambulodomus* | Pale Brown | Namibia | -24.76715 | 13.8518 | 200 - 300 | 7/10/24 |
| NAM62b | *Suberites ambulodomus* | Pale Brown | Namibia | -24.7204 | 13.8731 | 200 - 300 | 8/10/24 |
| NAM63a | *Suberites dandelenae* | Pale Brown | Namibia | -24.6992 | 13.88535 | 200 - 300 | 8/10/24 |
| NAM66d | *Suberites dandelenae* | Pale Brown | Namibia | -24.7059 | 13.8268 | 200 - 300 | 8/10/24 |
| NAM74d | *Suberites ambulodomus* | Pale Brown | Namibia | -24.23335 | 13.8472 | 200 - 300 | 9/10/24 |
| NAM84d | *Suberites dandelenae* | Pale Brown | Namibia | -24.517 | 13.85595 | 200 - 300 | 10/10/24 |
| VIGB | *Suberites ficus/latus* | Orange | Galicia | 42.252 | -8.745 | 5 | 22/9/24 |
| VIGC | *Suberites ficus/latus* | Orange | Galicia | 42.252 | -8.745 | 5 | 22/9/24 |
| VIGD | *Suberites ficus/latus* | Orange | Galicia | 42.252 | -8.745 | 5 | 22/9/24 |
| VIGE | *Suberites ficus/latus* | Orange | Galicia | 42.252 | -8.745 | 5 | 22/9/24 |
| VIGF | *Suberites ficus/latus* | Orange | Galicia | 42.252 | -8.745 | 5 | 22/9/24 |
| VIGG | *Suberites ficus/latus* | Orange | Galicia | 42.252 | -8.745 | 5 | 22/9/24 |
| VIGH | *Suberites ficus/latus* | Orange | Galicia | 42.252 | -8.745 | 5 | 22/9/24 |
| POR23-175 | *Suberites ficus/latus* | Grey | Ireland | 52.6098 | -13.8983 | 279 | 6/10/23 |
| POR23-176 | *Suberites ficus/latus* | Grey | Ireland | 52.6098 | -13.8983 | 279 | 6/10/23 |
| POR23-93 | *Suberites ficus/latus* | Grey | Ireland | 53.445 | -12.5895 | 311 | 30/9/23 |
| CRO1 | *Suberites domuncula* | Orange | Croatia | 43.351111 | 15.177778 | 15 | 15/9/17 |
| CRO6 | *Suberites domuncula* | Orange | Croatia | 43.351111 | 15.177778 | 15 | 15/9/17 |

The sponges belonging to *Suberites* sp. nov. exhibited different color morphs, varying in external coloration, including gray, yellow, orange, red, brown, and blue. Samples from Ireland, Namibia, and the Balearic Islands (MEDITS-2024 survey) were collected by trawling, whereas those from Croatia and Galicia were collected by scuba diving. All samples were fixed in 96% ethanol immediately after collection. After 24 hours, the ethanol was replaced to ensure proper fixation of the specimens. When possible, photographs of living individuals were taken to record their coloration (Figure 2).

**Figure 2.**
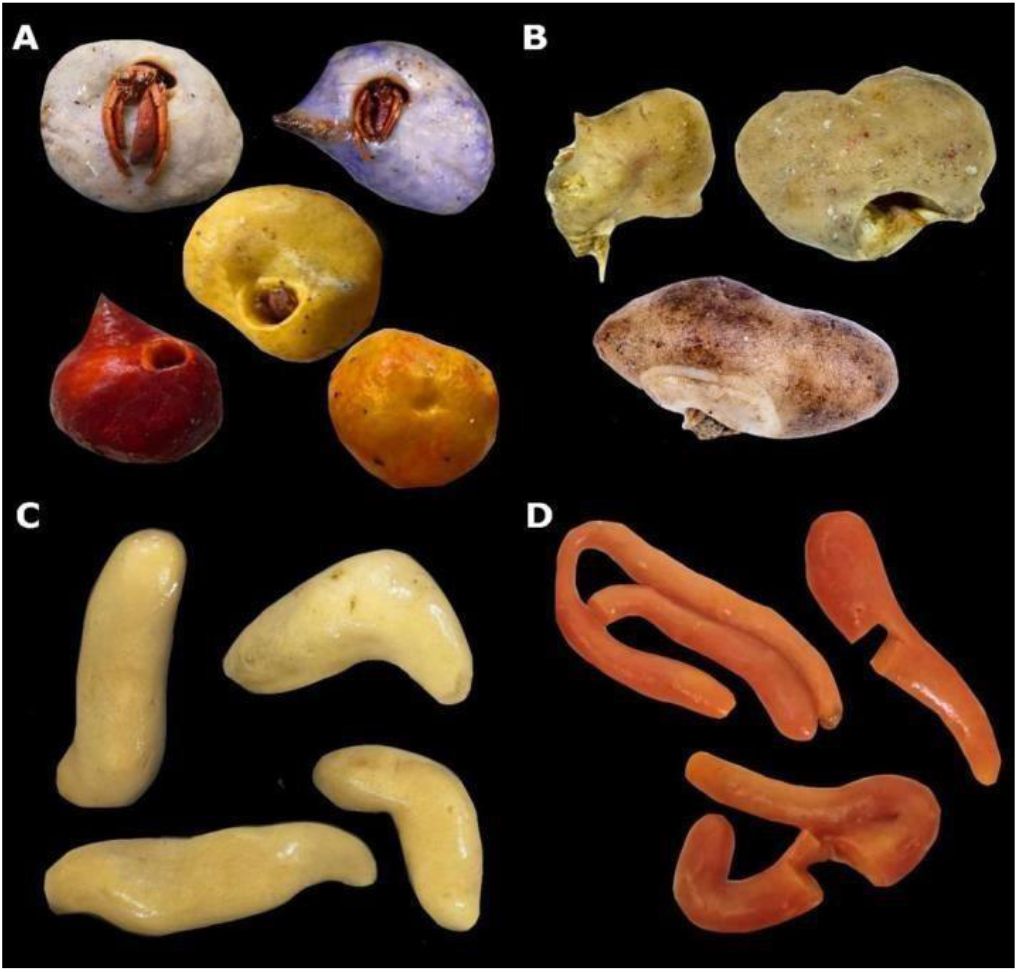
Photographs of representative samples used in the present study. A. Samples from *Suberites* sp. nov. from the MEDITS survey, consisting of specimens from the Balearic Islands from gray, yellow, orange, red, and blue color morphs. B. Samples of *Suberites ficus/pagurorum* from Porcupine Bank, Ireland. C. Samples of *Suberites ambulodomus* from Namibia. D. Samples of *Suberites ficus* from Vigo, Spain.

### Morphological analysis

For the morphological analysis of the spicules, a small fragment of each sponge, including part of the choanoderm and mesohyl, was excised. The tissue was immersed in a 100% solution of sodium hypochlorite at room temperature overnight to digest the organic matter. Subsequently, two washes with water and two with 96% ethanol were performed. The resulting solution was deposited onto a microscope slide, dried, and permanent preparations were made using DPX mounting medium. Spicules were visualized using an Olympus BX43 light microscope, and images were acquired with a UC50 camera using a 40× objective. The taxonomic identification of each individual was performed following specialized identification guides (Uriz 1986; Hooper & van Soest 2002) and bibliographic references (Olivi 1792; Johnston 1842; Solé-Cava & Thorpe 1986; Samai et al., 2017; Payne et al., 2024), based on spicule types and sizes. Spicule measurements were performed on a pair of individuals from each sampling site. Additionally, one individual of each of the differentiated color morphs from the MEDITS samples of *Suberites* sp. nov. was measured. For each sample, 10 spicules per category were measured, recording the length and width of each spicule. Data analysis and visualization were performed in R (R Core Team 2020) using the *ggplot2* package (Wickham 2016).

### DNA extraction, amplification, and sequencing

DNA was extracted from a piece of ca. 1 cm^3^ of all 43 samples analyzed using the Speertools Tissue DNA extraction kit (BIOTOOLS B&M Labs, S.A.), following the manufacturer’s recommendations, except for the cell lysis time, which was extended overnight. The quantity and quality of the extracted DNA were assessed using a NanoDropTM spectrophotometer. For most samples, a mitochondrial fragment of cytochrome c oxidase subunit I (COI) was amplified using the primers HCO 1490 (5’-TAAACCTTCTGGATGTCCAAAAAATCA-3’) and LCO 2198 (5’-GGTCAACAATCATAAAGATATTGG-3’) (Folmer et al., 1994). In some cases (Appendix 1, Table A1.3), the LCO primer was used in combination with Cox1-R1 (5’-TGT TGR GGG AAA AAR GTT AAA TT-3’), which amplifies the “Folmer” region as well as the “co1-ext” region, which has been used in other sponge barcoding studies (Rot et al., 2006). The following PCR program was used to amplify these fragments: 95°C/5 min – (95°C/30 s – 58°C/30 s – 72°C/1 min) × 35 cycles – 72°C/10 min. For some samples, the annealing temperature was reduced to 54°C. The total reaction volume was 13 µl, including 10 µl of VWR® Red Taq DNA polymerase master mix (1.1×), 1 µl of *forward* primer, 1 µl of *reverse* primer, and 1 µl of extracted DNA from each sample. PCR products were subsequently verified on a 1.5% agarose gel at 90 V for 35 min to determine whether amplification had occurred. Amplified PCR products were purified using ExoSAP-IT. Sequencing was performed using the dideoxynucleotide chain-termination method (Sanger method) by Macrogen (https://dna.macrogen.com/#), using the primer pairs indicated above.

### Phylogenetic analysis and haplotype network

Individual sequence reads were assembled and trimmed to obtain consensus sequences using Geneious v.6.1.6 (Kearse et al., 2012) (https://www.geneious.com). A total of 43 GenBank sequences from several species of the genus *Suberites*, as well as from other closely related genera, were included in the phylogenetic analysis (Appendix 1, Table A1.2). Multiple sequence alignment, 658 bases in length, was performed using MAFFT v.7.309 (Katoh & Standley 2013) for 25 different taxa and a total of 86 sequences.

The evolutionary model was selected using the Akaike Information Criterion (AIC) in jModelTest (Posada 2009), with HKY FU+IU+G4m obtained as the model that best fit the alignment. Phylogenetic analysis was performed using the maximum likelihood method in RAxMLGUI v2.0.15 (Edler et al., 2021) with 100 bootstrap replicates. The resulting tree was visualized and edited using FigTree v1.4.4 (Rambaut 2018), and was also edited using Adobe Illustrator (Adobe Systems Inc.). Unique haplotypes were identified from the sequence alignment, and a haplotype network was constructed using PopART v1.7 (Leigh & Bryant 2015) with the median-joining network method (Bandelt et al., 1999).

### Microbial composition and structure

In the 25 specimens of *Suberites* sp. nov. from the MEDITS campaign exhibiting color variation, the hypervariable V4 region of the 16S rRNA gene was amplified to test whether the microbiome composition is correlated with the colour. For this purpose, the primers 515 FY (Parada et al., 2016) and 806 R (Apprill et al., 2015) were used. A dual-indexing strategy was used to label each sample, as described by Kozich *et al.,* (2013). These primers contain degenerate bases to avoid the previous bias against Crenarchaeota/Thaumarchaeota and the SAR11 clade of Alphaproteobacteria. PCRBIO HiFi polymerase (PCR Biosystems Ltd) was used under the following PCR conditions: 95°C/5 min – (95°C/30 s – 58°C/30 s – 72°C/40 s) × 28 cycles – 72°C/5 min. DNA amplification was performed in duplicate, and PCR products were analyzed on a 1.5% agarose gel to determine amplification success and band intensity. PCR products were purified using Agencourt AMPure XP magnetic beads (Beckman Coulter Inc.), and libraries were prepared using the Nextera XT DNA Library Preparation Kit (Illumina Inc.). Next-generation sequencing was performed at NHMUK using an Illumina MiSeq with v3 chemistry (2 × 300 bp). Raw amplicon sequence reads were deposited in the Sequence Read Archive (SRA) under BioProject accession number PRJNA970968.

Sequences from the 16S V4 region were processed using the UPARSE processing pipeline (48). First, *usearch* was used to merge paired-end sequences, allowing a maximum of 5 base differences (*-fastq_mergepairs, -fastq_maxdiffs 5*), and primer sequences were removed (*-fastx_truncate*). The *vsearch* tool was used to perform quality control of the consensus sequences (*--fastq_stats*) and to filter them (*--fastq_filter*), retaining 92.76% of the sequences, which had a length of 252 bp. These sequences were clustered by identity, and unique sequences were identified while calculating the abundance of each sequence (*-fastx_uniques*). Amplicon denoising was performed using the UNOISE3 algorithm (Edgar 2013) to remove chimeras, sequencing-error reads, PhiX sequences (used as a sequencing control), and low-complexity sequences potentially caused by Illumina sequencing artifacts. Zero-radius operational taxonomic units (ZOTUs) were generated using sequences with 100% identity. ZOTU alignment and taxonomic assignment were performed using SINA v1.2.11 (Quast et al., 2013), with the SILVA 138_2 database. Finally, sequences with low alignment quality (<75%) or those that could not be classified as bacteria or archaea were removed from the analysis, as were sequences identified as mitochondria or chloroplasts.

Data analysis and visualization were performed in R (R team Core) using the *vegan v2.5-6* (Oksanen et al., 2015) and *ggplot2* (Wickham 2016) packages. To visualize the relative abundances of the different taxonomic groups at the class level, a stacked bar plot was generated, with samples separated according to color morph (gray, yellow, orange, red, brown, and blue). Subsequently, to evaluate the ordination of the samples, a Bray–Curtis dissimilarity distance matrix was calculated based on the relative abundance table of ZOTUs transformed using log2. These analyses allow patterns in microbial community composition to be visualized. A principal coordinates analysis (PCoA) was performed to visualize this distance matrix, with each color morph represented by a different color. To assess whether sponge color morph was the factor determining microbial composition, homogeneity of variances was first assessed using the *permutest* dispersion test implemented in the *vegan* package, followed by a non-parametric analysis of variance (PERMANOVA) using the *adonis* function in R. Finally, core communities (ZOTUs present in all samples belonging to the same group) and ZOTUs exclusive to each color morph were identified. The results of this analysis were visualized using an UpSet plot with the R package *UpSetR* (Conway et al., 2017).

## RESULTS

### Morphological analysis

The morphological study of spicules for the identification of *Suberites* species focused on the composition and size of the different spicule types, following the “(minimum measurement-(mean)-maximum measurement)” format for both length and width (Appendix 1, Table A1.1). The MEDITS samples exhibited styles and oxeas (Figure 3A), with styles having one rounded end and oxeas having both ends pointed (Figure 4A).

**Figure 3.**
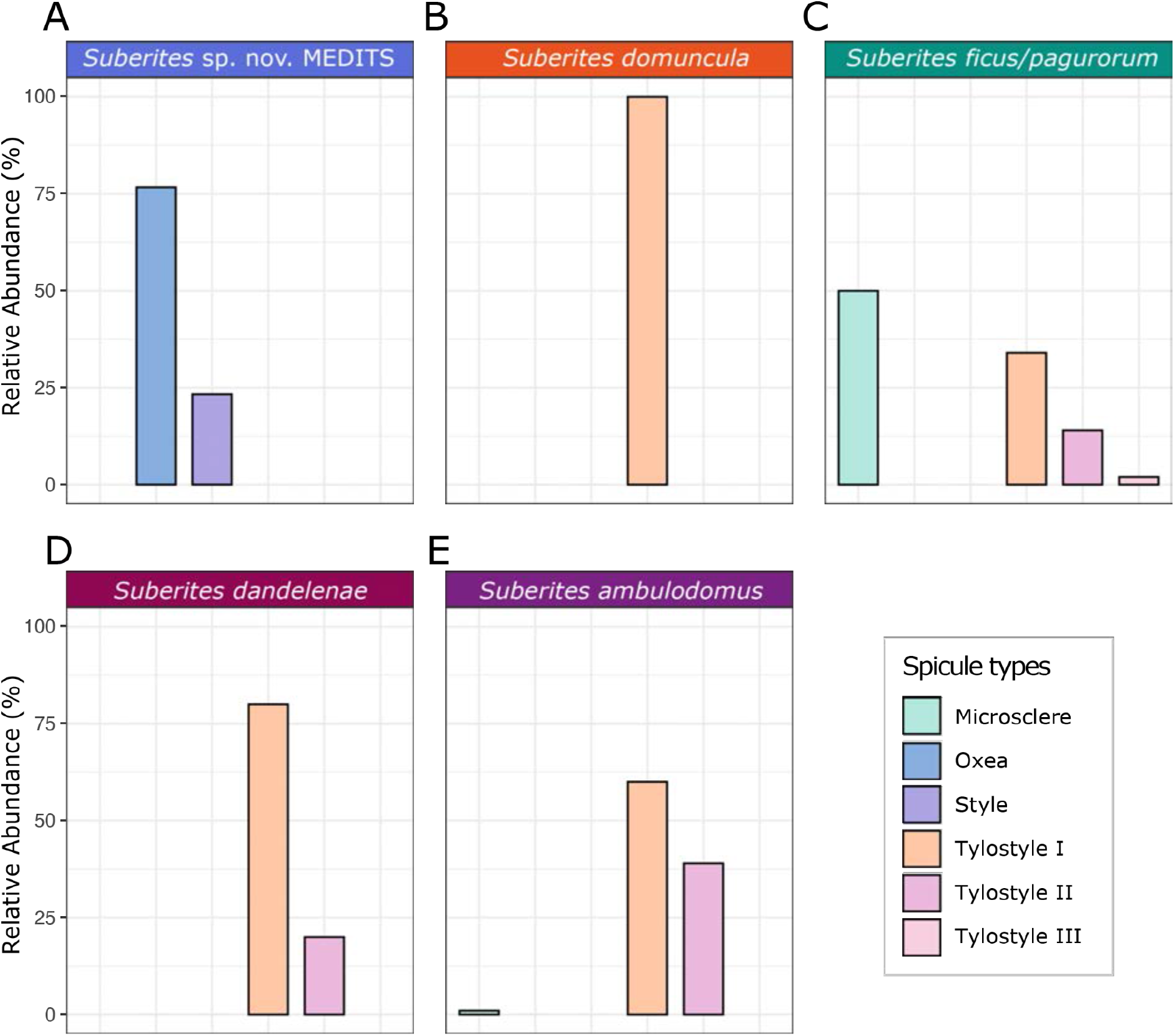
Bar plots representing spicule composition based on the relative abundances of the different spicule types. A. *Suberites* sp. nov. MEDITS. B. *Suberites domuncula*. C. *Suberites ficus/pagurorum*. D. *Suberites dandelenae*. E. *Suberites ambulodomus*.

**Figure 4.**
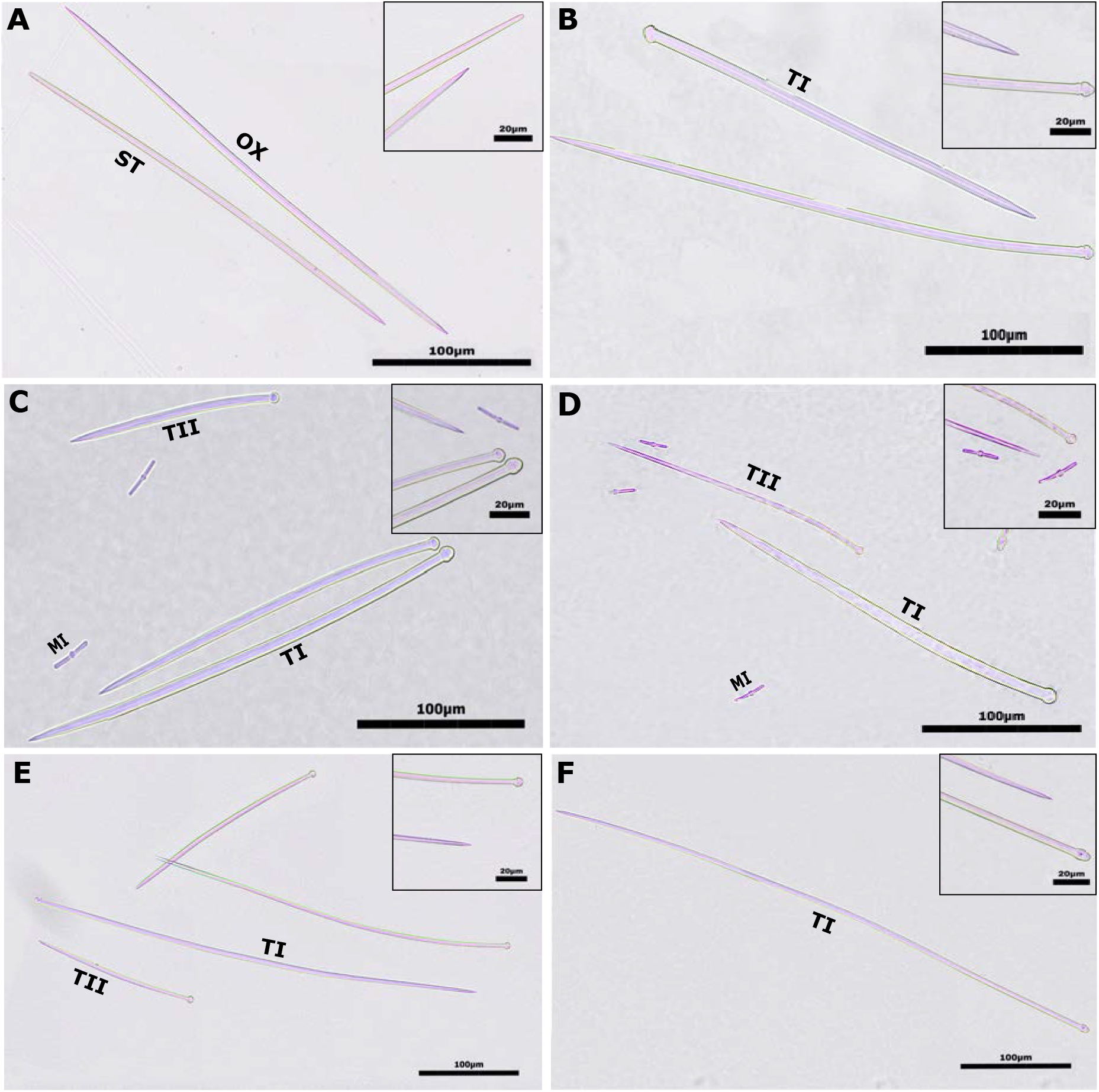
Micrographs of representative spicule types of each species. The small insets in each photograph show the ends of the spicules. Abbreviations correspond to the spicule type: OX, oxeae; ST, style; TI, larger tylostyle category; TII, smaller tylostyle category; MI, microsclere. A. *Suberites* sp. nov. from MEDITS. B. *S. domuncula*. C. *S. ficus* from Galicia. D. *S. ficus* from Ireland. E. *S. ambulodomus*. F. *S. dandelenae*.

The dimensions of the styles were 140-(240)-340 μm × 3-(4.5)-6 μm, whereas those of the oxeas were 173-(300.5)-387 μm × 2-(4)-6 μm, with no distinctions made among the different color morphs of the species. The *S. domuncula* samples from Croatia exhibited only tylostyles measuring 190-(317)-398 μm × 3-(4.5)-6 μm (Figures 5B and 6B). The spicules of *S. ficus/pagurorum* from Galicia consisted of tylostyles and microscleres (Figures 5C and 6C), with two classes of tylostyles measuring 228-(304)-365 μm × 5-(6.3)-8 μm and 122-(158.7)-232 μm × 3-(5.3)-8 μm, and microscleres measuring 18-(25.7)-33 μm × 1-(2)-2 μm. In one individual, a third category of notably thinner tylostyles was observed, measuring 190-(256)-315 μm × 2-(3)-4 μm. The *S. ficus/pagurorum* samples from Ireland exhibited tylostyles and microscleres (Figures 5C and 6D). The tylostyles comprised two categories, one measuring 271-(335.5)-419 μm × 7-(9)-11 μm and the other 145-(210.5)-281 μm × 5-(7.5)-9 μm, while the microscleres measured 17-(24)-33 μm. In one of the samples, a third category of tylostyles was observed, as in one of the samples from Galicia, measuring 139-(198)-250 μm × 3-(4)-5 μm. The *S. ambulodomus* samples from Namibia shared the same spicule composition, as they contained only tylostyles (Figure 3E); however, two categories of this spicule type were present, one measuring 310-(460.5)-575 μm × 3-(4.5)-6 and the other 117-(153.5)-2050 (Figure 4E). In contrast, *S. dandelenae* exhibited only one category of tylostyles (Figures 5D and 6F), with tylostyles measuring 482-(568)-643 μm × 4-(5)-6 μm.

**Figure 5.**
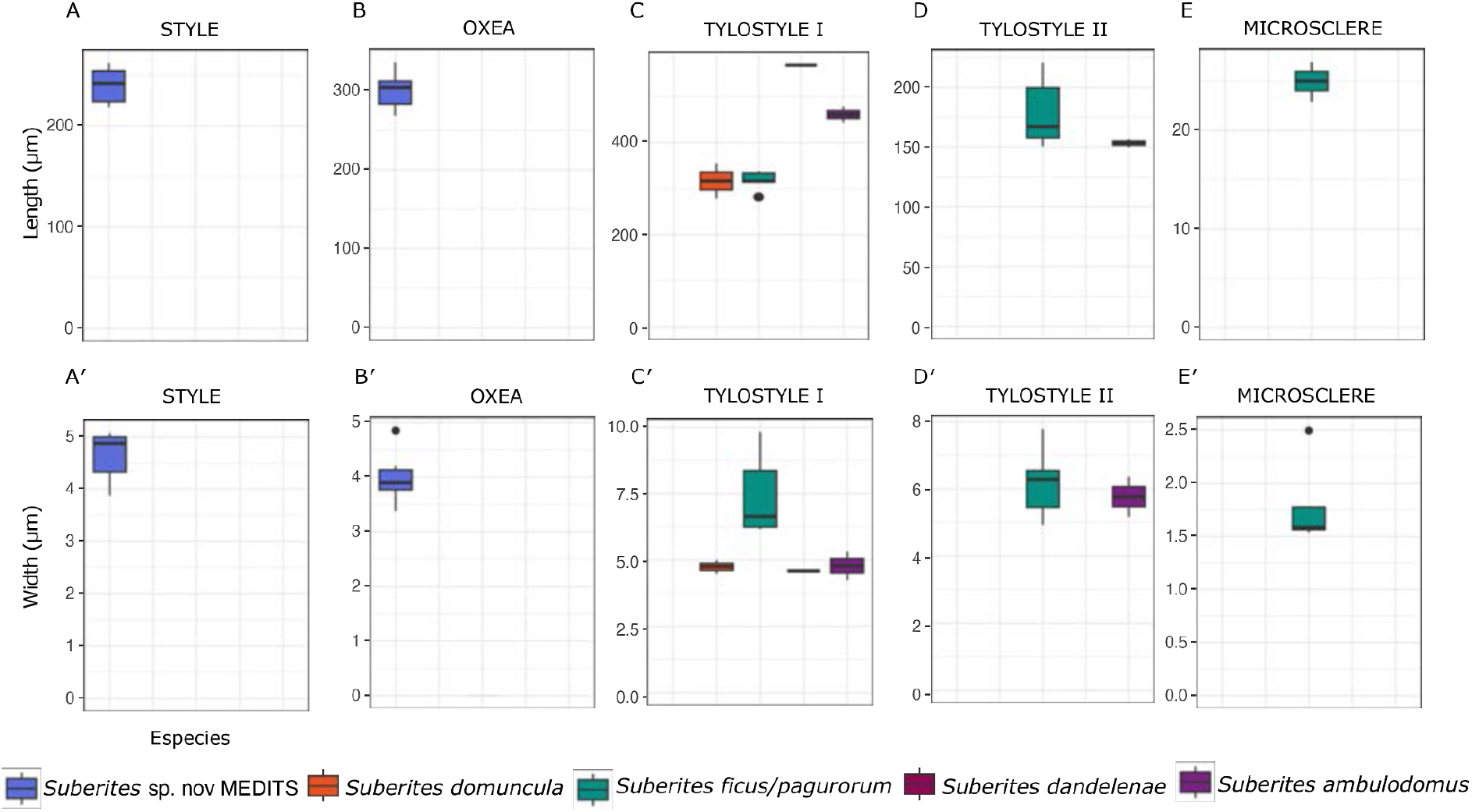
Box plots representing the mean spicule measurements across all species according to each spicule type. The upper plots show length measurements, whereas the lower plots show width measurements. Horizontal lines indicate the mean, vertical lines indicate the minimum and maximum values, and points represent outliers. A and A′. Styles. B and B′. Oxeas. C and C′. Large tylostyles. D and D′. Small tylostyles. E and E′. Microscleres.

**Figure 6.**
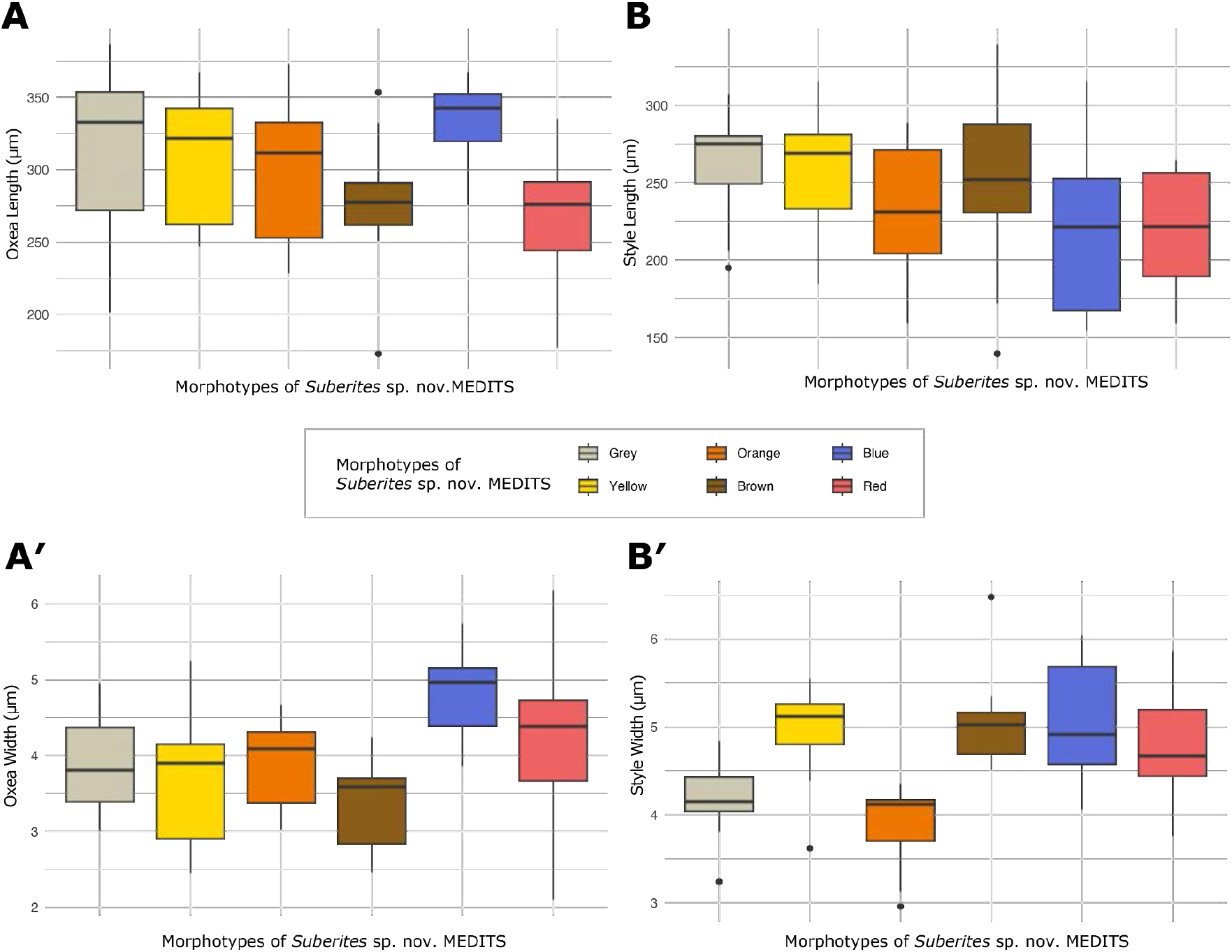
Box plots representing the measurements obtained from the spicules of *Suberites* sp. nov. MEDITS, from a single individual of each color morph. Horizontal lines indicate the means, vertical lines indicate the minimum and maximum values, and points represent outliers. A. Oxeas length. A′. Oxeas width. B. Styles length. B′. Styles width.

The main differences among species were related to the spicule types present in each species, as shown in Figure 3. Oxeas and styles were exclusive to *Suberites* sp. nov. MEDITS (Figure 3A), whereas microscleres were only present in *S. ficus/pagurorum* from Galicia and Ireland (Figure 3C). In contrast, the presence of tylostyles was not exclusive to a single group, although these differed among species, as type I tylostyles (the larger category) of *S. ambulodomus* and *S. dandelenae* from Namibia were the longest (Figure 5C), but not the widest, with those of *S. ficus/pagurorum* showing the greatest width (Figure 5C). Among the smaller tylostyles (type II tylostyles), those of *S. ambulodomus* were slightly shorter than those of *S. ficus/pagurorum* (Figure 5D), but no major differences in width were observed either (Figure 5D′).

To determine whether spicule differences existed among the color morphs present in *Suberites* sp. nov. MEDITS, additional measurements were performed separately for each color morph. The gray morph had styles measuring 195-(262)-307 × 3-(4)-5 μm and oxeas measuring 202-(313)-386 × 3-(4)-5 μm; the yellow morph had styles measuring 185-(255)-315 × 4-(5)-6 μm and oxeas measuring 247-(307)-368 × 2-(4)-5 μm; the red morph had styles measuring 159-(218)-265) × 4-(5)-6 μm and oxeas measuring 177-(268)-335 × 2-(4)-6 μm; the orange morph had styles measuring 159-(234)-288 × 3-(4)-4 μm and oxeas measuring 228-(301)-373 × 3-(4)-5 μm; the brown morph had styles measuring 140-(250)-340 × 5-(5)-7 μm and oxeas measuring 173-(271)-353 × 2-(3)-4 μm; and the blue morph had styles measuring 154-(220)-315 × 4-(5)-6 μm and oxeas measuring 276-(336)-368 × 4-(5)-6 μm.

Figure 6 graphically represents the measurements separated by color morph, showing some degree of variability. The blue morph had the largest oxeas (both longer and wider), whereas the brown morph had thinner oxeas that were similar in length to those of the red morph. The gray morph also had the longest styles, although they were thin, nearly as thin as those of the orange morph. Notably, the red morph had the shortest styles and oxeas, although both spicule types were among the widest. Nevertheless, no notable differences were observed among the color morphs.

### Phylogenetic analysis and haplotype network of *Suberites* spp

The 43 specimens analyzed in our study belonged to five different species, one of which potentially represents a new species. Each species occurred in a different sampling site, except for *S. ambulodomus* and *S. dandelenae*, both of which were collected from Namibia (Figure 7A).

**Figure 7.**
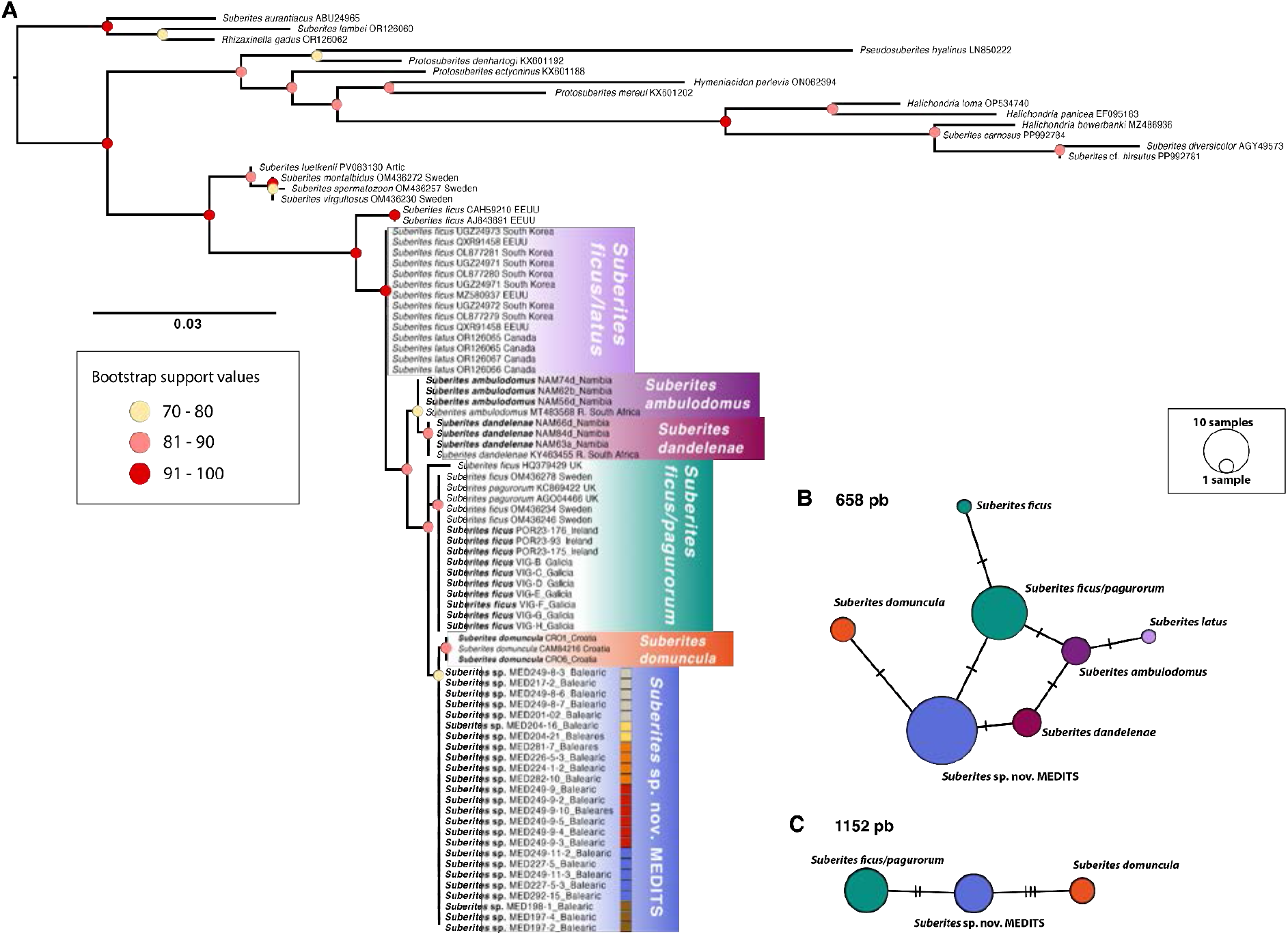
A. Maximum-likelihood phylogenetic tree of the *cox1* locus. The GenBank accession numbers and their locations (when known) are shown following the names; sequences included in this study are highlighted in bold. Colored squares indicate sponge coloration for *Suberites* sp. nov. MEDITS. Support values are indicated by circles at each node, associated with different colors: 70–80 (yellow), 81–90 (pink), and 91–100 (red). The scale bar indicates substitutions per site. B. COI haplotype network. Fragment of 658 base pairs for the different *Suberites* species. Circle size is proportional to the number of individuals represented by each haplotype, and the lines between them indicate the number of mutations separating them. C. Haplotype network of the 1152-base-pair COI fragment.

The maximum-likelihood tree recovered the genus *Suberites* as paraphyletic, as members of this genus were distributed across several clades, some of which also contained species from other genera (Figure 7A). The sequences generated in our study clustered in a moderately supported clade and formed a sister clade to sequences of *Suberites latus* from Canada and *Suberites ficus* from the United States and South Korea. Among the samples analyzed in this study, two major sister clades with moderate support values were distinguished.

On the one hand, one clade grouped the samples collected in Namibia into two subclades belonging to two different species: *S. ambulodomus* and *S. dandelenae*. Notably, the sequences of these two species were identical to sequences of the same species previously deposited in GenBank from South Africa. On the other hand, another clade grouped the samples with an Atlantic-Mediterranean distribution. This clade included one individual of *S. ficus* from the United Kingdom, a clade referred to as *S. ficus/pagurorum* (grouping samples of *S. pagurorum* and *S. ficus* from the North Atlantic), a clade containing individuals of *S. domuncula* from Croatia (two of which were sequenced in this study), and a clade designated *Suberites* sp. nov. MEDITS, which grouped all samples collected from the Balearic Islands.

To obtain a more visual representation of the genetic distances among the *Suberites* species present in the phylogenetic tree, haplotype networks were constructed using COI alignments of different lengths (Figures 7B and 7C). In Figure 7B, based on the 658-bp fragment, a clear separation was observed among the haplotypes of the different species, with only a single base difference between them. Each species appeared as a single haplotype; that is, there were no sequence differences within each species. The *S. dandelenae* and *S. ambulodomus* samples from Namibia differed by only a single base. There was also a single base difference between the *Suberites* sp. nov. MEDITS samples and *S. domuncula* and *S. ficus/pagurorum*. In turn, the *S. ficus* sample from the United Kingdom exhibited one mutation relative to *S. ficus/pagurorum*, whereas the *S. latus* sample from GenBank (OR126065) showed a haplotype close to that of *S. ambulodomus*, also separated by a single mutational step. The haplotype network in Figure 7C was constructed using a 1152-base alignment including only specimens of *S. domuncula*, *S. ficus/pagurorum*, and *Suberites* sp. nov. MEDITS detailed in Appendix 1, Table A1.3. In this case, a clear separation between species was also observed, with a single haplotype for each species, and we detected a distance of two and three mutations between *S. ficus/pagurorum* and *S. domuncula* and *Suberites* sp. nov. MEDITS, respectively. It should be noted that only some representative samples of *Suberites* sp. nov. MEDITS were included in this haplotype network (Figure 7C), as indicated in Appendix 1, Table A1.3.

### Microbiome analysis of *Suberites* sp. nov. MEDITS

The microbial composition plot (class-level taxonomy) shown in Figure 8 indicated that all color morphs of *Suberites* sp. nov. MEDITS were highly similar to one another.

**Figure 8.**
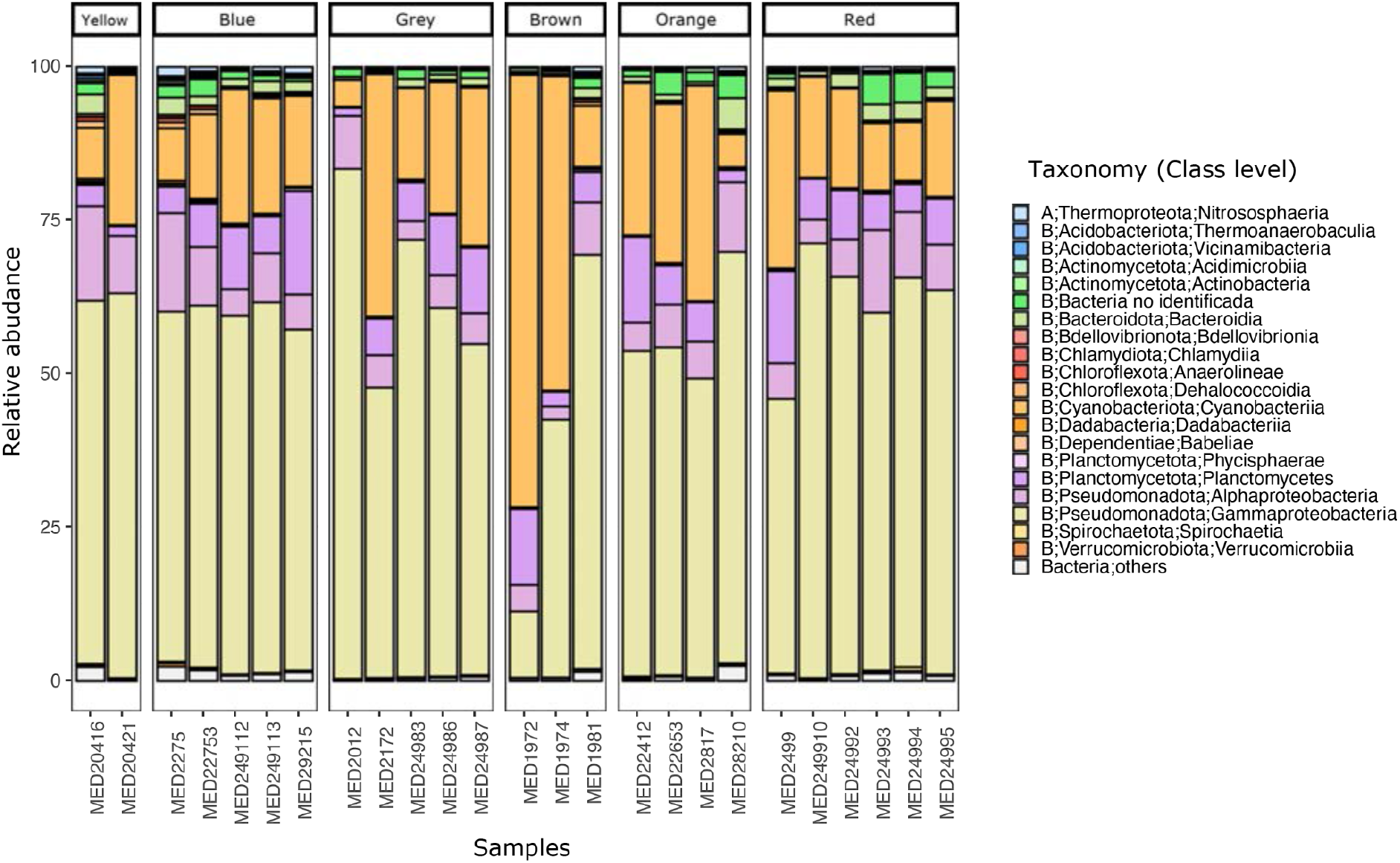
Stacked bar plot of the microbial composition of *Suberites* sp. nov. MEDITS sponges, separated by color morph. Taxonomic groups are identified at the class level, and those with abundances <10% are classified as “Bacteria; other”.

Among the prevalent microbial groups, the phylum Pseudomonadota was particularly prominent, generally representing between 50–75% of the microbiome of each sponge. Within this phylum, the class Gammaproteobacteria was the dominant class. The second most abundant group was the class Cyanobacteriia, whose relative abundance was generally between 20–25%. The presence of Planctomycetes and Bacteroidia was also notable, with relative abundances of approximately 10–15% and 3–5%, respectively. Specifically, in samples MED1972 and MED1974 (samples of the brown color morph; Table 1), the relative abundances of the bacterial groups did not follow the pattern observed in the remaining specimens analyzed, and higher values of cyanobacteria were observed (approximately 25%).

The principal coordinates analysis (PCoA) shown in Figure 9 indicated that the two main axes explained 50.4% of the variance in microbial composition (specifically, PCoA1 explained 31.6% and PCoA2 explained 18.8%). The ordination shown in this PCoA revealed significant separation among color morphs according to the PERMANOVA results (R² = 0.28, p-value < 0.05), with the “color” factor explaining 28% of the variance in the microbiome of these samples (Appendix 2, Tables A2.1 and A2.2).

**Figure 9.**
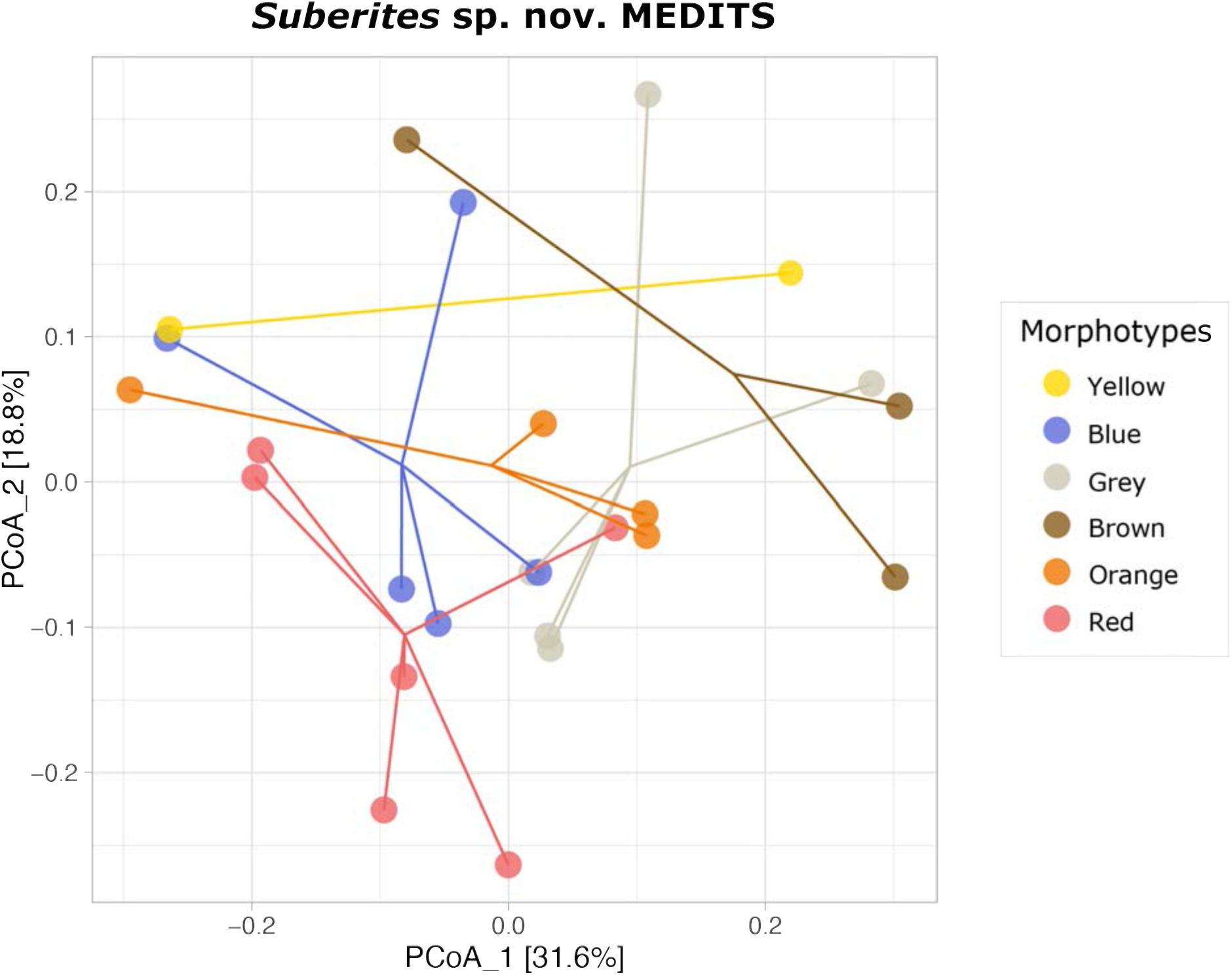
Principal coordinates analysis (PCoA) showing differences based on Bray–Curtis beta diversity in the microbial composition of the different color morphs of *Suberites* sp. nov. MEDITS. Colors correspond to the sponge color morphs.

Finally, an UpSet plot was generated to visualize how many ZOTUs were shared among and how many were exclusive to each of the color morphs (Figure 10). The total number of ZOTUs present in all samples was 221, while the number present in each color morph ranged from 240 (blue) to more than 400 (yellow), values resulting from the sum of shared and exclusive ZOTUs. Regarding ZOTUs exclusive to each color, values ranged from 196 for the yellow morph to 13 for the blue morph; all are available in Appendix 3 (Tables A3.1–6). Among these, several bacterial groups capable of producing pigments were identified (Appendix 3, Tables A3.1–6): in the gray morph, *Synechococcus* CC9902, *Pseudoalteromonas*, *Marinobacter*, *Maricinella*, and *Ruegeria*; in the yellow morph, *Fluviicola*, *Pelagibius*, *Salegentibacter*, and *Spirochaeta*; in the orange morph, *Pseudoalteromonas* and *Bremella*; in the red morph, Pseudomonadaceae, Cyanobiaceae, *Arenibacter*, *Jeotgalicoccus*, *Alteromonas*, *Marinobacter*, and Paracoccaceae; in the brown morph, Cellvibrionaceae, *Synechococcus* CC9902, *Cyanobium*, *Acinetobacter*, *Maricinella*, and *Flavobacterium*; and in the blue morph, *Synechococcus* CC9902, *Luminiphilus*, *Salinicoccus*, and Phycisphaeraceae.

**Figure 10.**
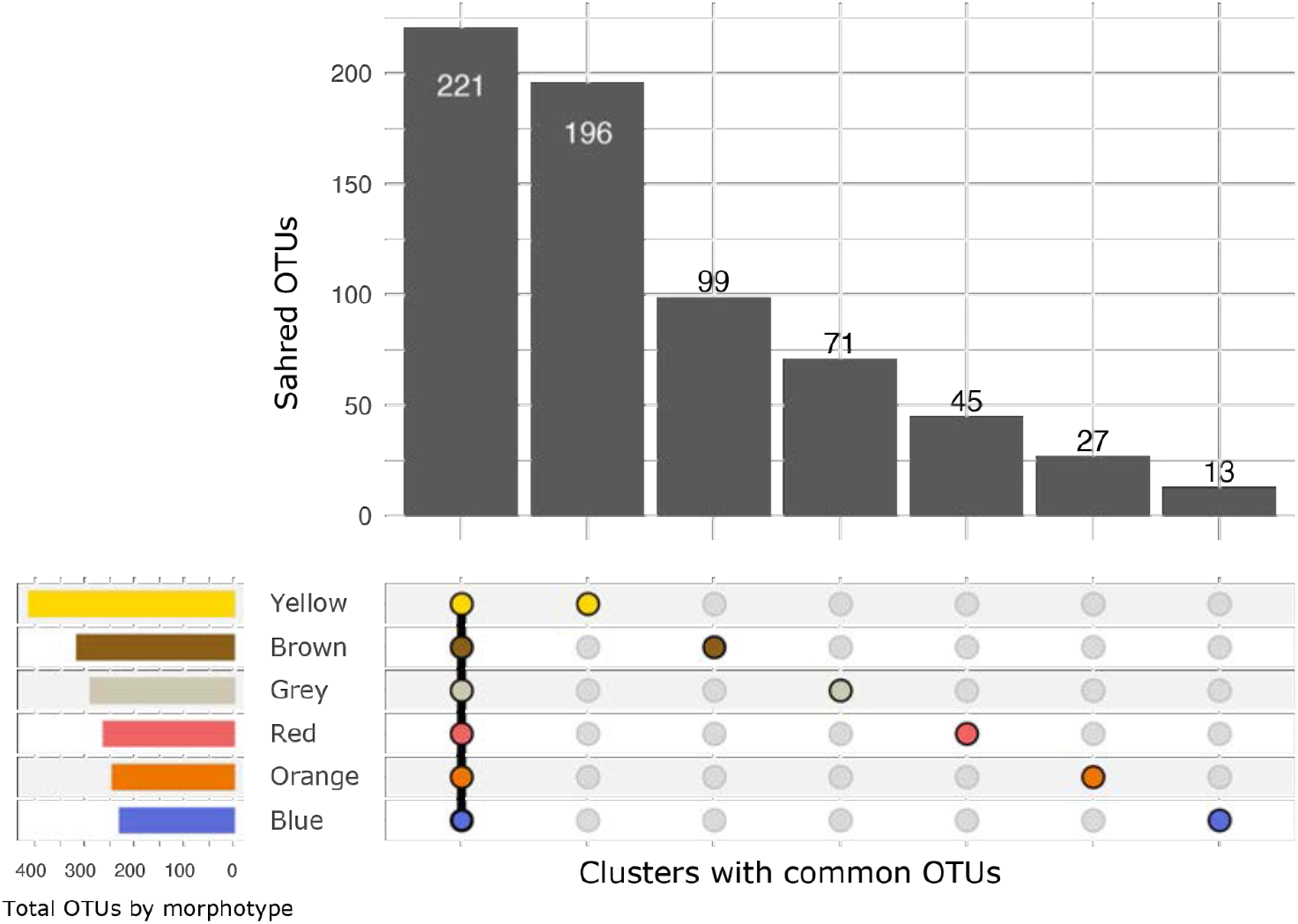
UpSet plot showing the ZOTUs shared among all color morphs and those exclusive to each morph. Colors correspond to the sponge color morphs. Points indicate which sponge group contains a given number of ZOTUs, with the first column indicating the ZOTUs shared by all color morphs.

## DISCUSSION

The phylogenetic analyses placed all the individuals of *Suberites* sp. nov. MEDITS studied within the same species, confirming that, at the genetic level, there are no differences despite the variations in coloration. Furthermore, all samples from the Mediterranean Sea may correspond to a new *Suberites* species that has so far only been found in the Balearic Islands. This species shows clear differences in spicule morphology compared with its sister species, *S. domuncula*. Analyses of the microbial composition of these samples revealed significant differences in the microbiome composition of individuals with different colorations. In addition, certain ZOTUs exclusive to each color morph were identified that produce pigments that could be related to the specific colorations observed in the sponges.

### Morphological study

External and internal morphology is key to the taxonomic identification of sponges. In most studies, external characteristics are the first features used to make a species-level identification (Hooper & van Soest 2002; Fortunato et al., 2020). In addition to sponge size, characteristics such as color, consistency, morphology, and the morphology of the oscula are carefully examined. The MEDITS samples showed external characteristics very similar to those of *S. domuncula*; the globular morphology and different color morphs were features shared by both species. However, this was not the case for the spicule morphology. While *S. domuncula* presented exclusively tylostyles (although, according to the literature, it may also present some oxeas and styles (Hartman 1958; Uriz 1986), the MEDITS samples corresponding only possessed oxeas and styles, with no tylostyles in their skeletal composition (Figures 5A and 5B; Figures 6A and 6B). The measurements of the tylostyles from the *S. domuncula* individuals analyzed fell within the range of measurements reported by Fortunato *et al.,* (2020) for tylostyles of this species, but were larger than those reported by Hartman (1958) and smaller than those reported by Müller *et al.,* (2006). The different color morphs exhibited by *Suberites* sp. nov. MEDITS were very similar in their spicule composition, suggesting that, despite their different coloration, they belong to a single species. All morphs presented the same spicule types (only oxeas and styles), with slight variations in their length and width. However, the blue sponges stood out due to their greater spicular variation, presenting longer and wider oxeas and styles with greater variation in size and width (Figure 6), although more individuals of this morph should be measured to determine whether these differences are significant.

Externally, the samples belonging to *S. ficus/pagurorum* from Ireland and Galicia differed both in shape and color depending on the sampling site, being globose and beige in Ireland (Figure B), and elongated and orange in Galicia (Figure 3D). The presence of centrotylote strongyles, together with tylostyles, characterizes and defines this group and did not differ between the different populations. Its spicule composition corresponds to that reported for *S. ficus* in other studies (Hartman 1958; Solé-Cava & Thorpe 1986). However, the size of its tylostyles falls within the range reported for *S. pagurorum* by Solé-Cava & Thorpe (1986), *S. latus* by Turner *et al.,* (2024), and *S. ficus* (from France) by Hartman (1958). Therefore, the taxonomic assignment of this species (*S. ficus/pagurorum*) based on morphology is not conclusive, as the taxonomic boundaries between *S. ficus* and *S. pagurorum* have not been clearly defined. The two species from Namibia, *S. ambulodomus* and *S. dandelenae*, were very similar externally, with an elongated globose shape and beige coloration (Figure 3C). At the spicular level, both species presented type I tylostyles, longer in *S. dandelenae* (Figure 5C). The key difference lies in the presence of a second, smaller type of tylostyle in *S. ambulodomus* (Figure 4E). Nevertheless, the type I tylostyles of both species were larger than those measured in the study by Samaai *et al.,* (2017).

Spicule analysis proved to be essential for distinguishing the species examined in this study, as it revealed differences and similarities that are not evident from external morphology. The composition of spicule types was the most relevant spicular character for differentiating species in our study, whereas spicule size did not appear to play a relevant role in species separation. This is because the types of spicules present in a sponge are genetically determined, whereas their growth is linked to the availability of silicon in the environment (Uriz 2006). A previous study has detected even monthly and seasonal changes in spicule size depending on the time at which samples were collected (Jones 1991). Therefore, the use of spicule size as a character for separating species may not always have an evolutionary basis, but rather may be determined at the ecological level, making it necessary to use other types of analyses, such as genetic data.

### Genetic analysis

The phylogenetic analysis including the different species of the genus *Suberites* revealed that all *Suberites* sp. nov. MEDITS samples formed a single clade, which in turn was a sister clade to *S. domuncula* (Figure 7A). Notably, all *Suberites* sp. nov. MEDITS individuals exhibited exactly the same COI haplotype, with no genetic differences among the different color morphs. In the haplotype network constructed for the short COI fragment (Figure 7B), the MEDITS samples were separated by a single mutational step from *S. domuncula*, *S. ficus/pagurorum*, and *S. dandelenae*. When the haplotype network was analyzed using a longer COI fragment, including the most closely related species (Figure 7C), up to three base differences were detected. Considering that, in both sponges and cnidarians (particularly in some demosponges), the mutation rate of mitochondrial genes is much slower than in bilaterian animals (Lavrov 2016), the small differences observed in COI suggest that *Suberites* sp. nov. MEDITS represents a new species. This is further supported by the differences in spicule morphology discussed above.

As with the morphological analysis, the genetic distinction between *S. ficus* and *S. pagurorum* remains unresolved; for this reason, these samples are referred to as *S. ficus/pagurorum* in this study. The taxonomic dilemma surrounding these two species and their relationship with *S. domuncula* has also been observed in other studies using COI (Maldonado et al., 1999; Morrow et al., 2012) and in studies using the 28S marker (Morrow et al., 2012), which likewise failed to reach a consensus regarding their position in the phylogenies. In fact, according to Solé-Cava & Thorpe (1986), *S. pagurorum* is a subspecies of *S. ficus*. In addition, several *S. ficus* haplotypes have been distinguished depending on whether they originate from the Atlantic or Pacific Ocean, with the latter forming a clade with *S. latus*. Further genetic studies are needed, including, for example, complete mitochondrial genome sequencing or even low-coverage whole-genome sequencing, combined with morphological analyses using samples from both oceans, to shed light on these phylogenetic inconsistencies. Finally, *S. dandelenae* and *S. ambulodomus* from Namibia formed the sister group to the remaining clades containing *S. ficus/pagurorum*, *S. domuncula*, and *Suberites* sp. nov. MEDITS. This phylogenetic distribution was also observed in another study conducted by Payne *et al.,* (2024), although it differs from the results obtained by Turner *et al.,* (2024). In the latter study, the sister clade to the Namibian species was formed by *S. latus* and Pacific *S. ficus*.

### Ecological differences

The genetic analyses also revealed a close relationship between sampling locations and phylogenetic groupings. In most cases, clades corresponded to specific distribution areas that, in general, do not overlap. The distribution of *S. domuncula*, which was initially described in the Adriatic Sea (Olivi 1792), overlaps with that of *Suberites* sp. nov. MEDITS, as both occur in the Mediterranean Sea. Macroscopic identification of these two species may be confusing due to their similarities in external appearance and the fact that they commonly live in association with a hermit crab. Given the number of characteristics they share, it is possible that some specimens in previous studies were incorrectly identified as *S. domuncula* (Ferrer 1921), which could explain the inconsistencies observed in the literature regarding their spicules. *Suberites domuncula* and *Suberites* sp. nov. MEDITS should thus be considered as pseudo-cryptic species, since they are very similar externally, but small key differences can be observed at the spicular level. Future studies should analyze populations of both species to determine whether they coexist or, conversely, whether their distributions do not overlap. A detailed review of the literature would be essential to correct possible previous taxonomic misidentifications.

Among the remaining samples, it was also observed that those that were geographically closer were also genetically closer, that is, clades corresponded to sampling areas. However, no genetic differences were observed between the *S. ficus/pagurorum* samples from Galicia and Ireland despite their geographic separation, whereas the Namibian samples belonged to different species even though they were collected in the same area. Geographic distance is generally proportional to the degree of isolation (Blanquer & Uriz 2010), and therefore it is not unexpected that it may influence genetic distance to some extent.

### Influence of the microbiome on the coloration of *Suberites* sp. nov. MEDITS

Microbial composition in sponges has been studied in a large number of species (Erwin et al., 2012; Díez-vives & Riesgo 2024), and these microbial communities, which live in association with sponges, have been observed to exhibit species-specific relationships. This indicates that the main determinant of microbiome composition is sponge identity, with variations compared with other species, even when these are phylogenetically closely related (Thomas et al., 2016). Furthermore, this close relationship is maintained at the genotype level in some species, demonstrating the highly specific and finely tuned association between the microbiome and the sponge (Díez-Vives et al., 2020). In addition to the taxonomic identity of the sponge, microbial community composition may also be influenced by different environmental factors (Thomas et al., 2016) and may be subject to seasonal variation (Erwin et al., 2012).

As all *Suberites* sp. nov. MEDITS samples analyzed belong to the same species, our study focused on identifying microbiome differences associated with the distinct color morphs, with the results revealing significant variation in microbiome composition among them. In most of the samples analyzed, the dominant phylum was Pseudomonadota (Gammaproteobacteria and Alphaproteobacteria), as in many other sponge species (Thomas et al., 2016). Notably, this was not the case in two of the three samples of the brown morph, in which Cyanobacteria were dominant; this group was the second most abundant in the remaining samples (Figure 7). These bacterial groups enable an efficient organic matter cycle, favoring nutrient acquisition and recycling and contributing to the remineralization of organic matter (Graham & Tully 2021), and they have also been associated with the colored morphs of *P. ficiformis* (Burgsdorf et al., 2014). Another of the most abundant groups was Planctomycetes, which produce bioactive compounds that could help the sponge defend against other pathogenic microorganisms (Graça et al., 2016). These were not the only bacterial groups present in these samples, but they were the most abundant, representing approximately 90–95% of the total community in terms of relative abundance.

Significant differences were observed in microbial composition according to sponge coloration, although color was not the main factor explaining variation in the communities. In the PCoA (Figure 9), some clustering of individuals with the same coloration was detected, although other factors that were not evaluated could be explaining the observed differences. These differences were not apparent at the class taxonomic level, as shown in the bacterial composition bar plot, where all groups exhibited a similar composition (Figure 8). However, at the ZOTU level, differences among color groups were detected, with communities exclusive to each color morph being identified (Figure 10). When these exclusive ZOTU groups were analyzed, bacterial groups capable of producing pigments that may contribute to the coloration of these sponges were observed (Figure 9).

The yellow morph samples contained the highest number of exclusive ZOTUs, including bacteria from groups such as *Fluviicola*, *Pelagibius*, *Salegentibacter*, and *Spirochaeta*, which can produce pigments. These include carotenoids, pigments that are typically yellowish or orange in color (Breznak & 1969; Choi et al., 2009), similar to those found in the orange samples. The latter contained groups such as *Pseudoalteromonas* and *Bremella* (Godinho et al., 2021, Ren et al., 2024). In the red morph samples, bacterial groups such as *Alteromonas*, *Arenibacter*, Pseudomonadaceae, Cyanobiaceae, and *Jeotgalicoccus* were identified, which produce pigments that are red, pink, dark orange, green, and blue-green in color (Gerber & Gauthier 1979; Ivanova et al., 2001; Usman et al., 2017). In the gray morph samples, bacteria such as *Pseudoalteromonas*, *Maricinella*, and *Ruegeria* were found; among the pigments they produce are some that are gray, brown, and yellow in color (Romanenko et al., 2009; Ren et al., 2024). The brown morph contained bacteria that produce melanin, phycobiliproteins, and carotenoids, which are pigments with brown, orange, blue, and grayish colors. These are produced by bacteria belonging to the groups *Acinetobacter*, Cellvibrionaceae, *Cyanobium*, and *Flavobacterium* (Pagels et al., 2022; Raman et al., 2024). Finally, the blue morph also contained bacteria producing phycobiliproteins and bacteriochlorophyll *a*, which are blue, purple, and pink in color, associated with *Synechococcus*, *Luminiphilus*, Halieaceae, and Phycisphaeraceae (Fukunaga et al., 2009; Grébert et al., 2022), as well as carotenoids such as zeaxanthin, produced by *Salinicoccus* (Usman et al., 2018). The blue and orange morphs were the two groups with the lowest number of exclusive ZOTUs. This could be related to the fact that some sponges of the blue morph showed slight orange patches. It is possible that many of the ZOTUs present in the blue sponges also occur in the orange ones; therefore, when identifying ZOTUs exclusive to each color, many are shared and are not counted as exclusive, thereby reducing the representation of each group. This relationship is also reflected in the PCoA analysis, where the blue sponges cluster predominantly together with the orange sponges. In any case, further studies should be conducted to explore the ecological or evolutionary advantages (if any) that these colors may provide to the individuals in the habitats where they occur.

## CONCLUSION

In this study, intra- and interspecific differences at the genetic and morphological levels were determined among different species of the genus *Suberites*. This combined approach led to the discovery of a potentially new species in the Mediterranean Sea, *Suberites* sp. nov. MEDITS, which is pseudo-cryptic with respect to *Suberites domuncula*. Future studies will need to analyze additional samples of *Suberites* sp. nov. MEDITS from other parts of the Mediterranean and examine all of its morphological characteristics in greater depth in order to describe it properly. This new species exhibits a wide variety of color morphs (yellow, orange, red, blue, brown, and gray), which are not associated with detectable genetic differences in the COI marker or clear differences in spicule morphology, but rather appear to be mediated by the associated symbiotic bacteria. Each morphotype presents exclusive ZOTUs containing pigment-producing bacteria that may contribute to the characteristic coloration of each morphotype. These results support a role for the microbiome in determining, at least in part, the coloration of this sponge.

## APPENDIX 1

- **Table A1.1.** Spicule measurements in μm. Maximum measurement–(mean)–minimum measurement.

- **Table A1.2.** GenBank sequences used to construct the phylogenetic tree.

- **Table A1.3.** Samples included in the haplotype network of the 1,152-bp COI fragment.

## APPENDIX 2

- **Table A2.1.** Results of the dispersal analysis based on 999 free permutations.

- **Table A2.2.** Results of the PERMANOVA analysis based on Bray–Curtis dissimilarities, using 999 permutations, to assess the effect of colour on the microbial community composition of *Suberites* sp. nov. MEDITS.

## APPENDIX 3

- **Table A3.1.** Exclusive ZOTUs identified in red-morphotype samples of *Suberites* sp. nov. MEDITS.

- **Table A3.2.** Exclusive ZOTUs identified in grey-morphotype samples of *Suberites* sp. nov. MEDITS.

- **Table A3.3.** Exclusive ZOTUs identified in orange-morphotype samples of *Suberites* sp. nov. MEDITS.

- **Table A3.4.** Exclusive ZOTUs identified in brown-morphotype samples of *Suberites* sp. nov. MEDITS.

- **Table A3.5.** Exclusive ZOTUs identified in blue-morphotype samples of *Suberites* sp. nov. MEDITS.

- **Table A3.6.** Exclusive ZOTUs identified in yellow-morphotype samples of *Suberites* sp. nov. MEDITS.

